# Senescent cell survival relies on upregulation of lysosomal quality control mechanisms

**DOI:** 10.1101/2025.03.31.646397

**Authors:** Rachel Curnock, Bernadette Carroll

## Abstract

The accumulation of senescent cells is a key mediator of tissue and organismal ageing. Persistent activation of the growth regulator, mTORC1, even in the absence of growth factors and amino acids, supports senescence phenotypes, such as increased cell size and secretion of inflammatory factors. Here we extend this finding to show that senescence is associated with lysosomal accumulation of the low-density lipoprotein receptor (LDLR) and a failure of mTORC1 signalling to respond to changes in cholesterol. These observations are reflective of a broader dysfunction through the endo-lysosomal pathway, with Rab GTPases and phosphoinositides localised to atypical hybrid organelles. We propose that endosomal mistrafficking, in concert with increased autophagy and elevated lysosomal pH are contributing to an accumulation of undegraded material and lysosomal membrane damage. Our data indicate that in response to lysosomal dysfunction, senescent cells not only upregulate TFEB/TFE3-dependent lysosomal quality control but also upregulate lysosomal repair via PI4K2A-dependent PITT pathway. Perturbation of the lipid transport machinery required for lysosomal repair caused senescent cell death, revealing that targeting mechanisms of lysosomal repair has senolytic potential.

## Introduction

Cellular senescence is defined as an irreversible exit from the cell cycle. Senescent cells resist cell death and increase in cell size, mitochondrial mass and enhance the secretion of proinflammatory and pro-oxidant signals despite being non-proliferative. Gross changes in the contents and the morphology of the lysosome was one of the earliest defining hallmarks of senescence. For over a decade now, we have been aware of the importance of the autophagy-lysosome pathway to supply amino acids to maintain mTORC1 activity in senescence (Narita et al., 2011), which has since been shown to fuel the hypersecretory state in senescence known as the senescence-associated secretory phenotype (SASP) (Herranz et al., 2015; Laberge et al., 2015). We reported that mTORC1 is constitutively active in senescence, and unresponsive to changes in amino acids and growth factors (Carroll et al., 2017) and more recently we demonstrated that lysosomes in senescent cells are more susceptible to damage and TFEB-driven lysosomal quality control is essential for senescent cell viability (Curnock et al., 2023).

Central to regulating anabolism and catabolism, proliferating cells perceive nutrients to drive mTORC1 recruitment and activation at the lysosomal surface. How senescent cells bypass multiple signals and feedback loops to sustain mTORC1 signalling in the absence of stimuli is not fully understood. Lysosomal dysfunction has emerged as a unifying feature of many disease pathologies and is associated with normal ageing. We and others have shown that lysosomes in senescent cells are dysfunctional, characterised by elevated pH, evidence of membrane damage and diminished proteolytic activity. The lysosome fulfils an expanding repertoire of cellular functions and coordinates mTORC1 signalling through multiple mechanisms and in response an array of stimuli. Recent studies have elegantly demonstrated the role of the lysosome in detecting cholesterol and identified cholesterol sensing machinery that promote the recruitment of mTORC1 to lysosomes (Bayly-Jones et al., 2024; Castellano et al., 2017; Lim et al., 2019; Shin et al., 2022; Xiong et al., 2025; Zhao et al., 2025). Aberrant cholesterol metabolism is associated with many age-related diseases such as atherosclerosis, cancer, arthritis and diabetes, however, cholesterol sensing in senescence is unstudied.

An increasing number of lysosomal attributes are now known to influence mTORC1 activity including spatiotemporal positioning, phosphoinositide membrane composition and the fitness of the endosomal network that feeds into lysosomes as the terminal compartment. Hence, lysosomal dysfunction in senescence and ageing may have further reaching impacts on mTORC1 activity than we first appreciated. The importance of the lysosome to cellular homeostasis and ultimately cell survival is apparent by the evolutionarily conserved pathways that repair or degrade damaged lysosomes. The endosomal sorting complex required for transport (ESCRT) and autophagy machinery are vital multi-subunit protein complexes that mediate lysosomal repair and degradation, respectively (Maejima et al., 2013; Papadopoulos et al., 2017; Papadopoulos and Meyer, 2017; Radulovic et al., 2018; Skowyra et al., 2018). Alternative lysosomal repair pathways have recently been identified that rely on the rapid recruitment of the kinase, phosphatidylinositol-4 kinase type 2α (PI4K2A), to damaged lysosomes to drive cholesterol transfer-dependent repair (Radulovic et al., 2022; Tan and Finkel, 2022). The central kinase to this pathway, PI4K2A, also coordinates nutrient-controlled lipid conversion between PI(3)P and PI(4)P (Ebner et al., 2023). This lysosomal phosphoinositide heterogeneity underpins the functional switching of lysosomes between mTORC1-dependent anabolic growth signaling and the catabolic degradation of macromolecules. Thus, given the importance of the lysosome-autophagy pathway to sustained mTORC1 activity in senescence, understanding if these lysosomal attributes and repair mechanisms are affected in senescent cells could reveal novel senolytic targets.

Here, we show that mTORC1 signalling is refractory to cholesterol starvation in senescence and this is accompanied with an accumulation of the low-density lipoprotein receptor (LDLR) in the lysosome. We find that Rab GTPases and phosphoinositides are atypically localised in senescence and this could be promoting aberrant transmembrane protein delivery to the lysosome. We propose that endosomal mistrafficking in concert with elevated lysosomal pH is contributing to an accumulation of undegraded material leading to lysosomal membrane damage. We show that senescent cells are undergoing elevated levels of lysosomal repair with sustained PI4K2A lysosomal recruitment and concurrent increased lysosomal PI(4)P and cholesterol. We demonstrate that senescent cell viability relies on persistent lipid transfer-dependent lysosomal repair and we hypothesise that this is perturbing lysosomal phosphoinositide identity thus, impairing mTORC1 signalling in response to nutrient availability.

## Results and discussion

We previously found that persistent mTORC1 activity in the absence of growth factors and amino acids was a unifying feature of multiple models of senescence (Carroll et al., 2017), including the well-established oncogene-induced model of senescence (OIS, induced via 4OHT-inducible expression of HRasV12) we utilise in this current study. Cholesterol is essential for membrane biogenesis and vital for cellular growth. The mTORC1 pathway regulates *de novo* cholesterol synthesis and cholesterol uptake to meet the demands of the cell. It has recently emerged that cholesterol concentration at the lysosome is sensed and coupled to mTORC1-dependent anabolic signalling (Lim et al., 2019; Shin et al., 2022). Senescent cells have been reported to accumulate cholesterol in lysosomes and this has been linked to upregulation and mistrafficking of the cholesterol exporter ABCA1 (Roh et al., 2023). Dysregulation of cholesterol metabolism is associated with a plethora of age-associated pathologies. Therefore, we sought to examine whether cholesterol-dependent activation of mTORC1 is responsive in senescence.

At the lysosome, mTORC1 is responsive to discrete cholesterol pools; cholesterol transferred from the ER to the lysosomal limiting membrane (Lim et al., 2019) and cholesterol derived from low-density lipoprotein (LDL) which is trafficked to the lysosomal lumen where it stimulates Rag- and SLC38A9-dependent activation of mTORC1 (Castellano et al., 2017). In proliferating cells, following cellular cholesterol depletion with methyl-β cyclodextrin (MCD), supplementation with exogenous LDL or MCD-complexed cholesterol activated mTORC1 in a dose-dependent manner as measured by phosphorylation of canonical mTORC1 substrates (phospho S6 and ULK1) **(Fig. 1. A-D)**. In contrast, mTORC1 was unresponsive to exposure to either source of cholesterol in senescent cells **(Fig. 1. A-D)**. Using the cholesterol biosensor, mCherry-tagged D4H*, which exploits the cholesterol binding affinity of domain four from perfringolysin O (aka θ-toxin) produced by Clostridium perfringens (Lim et al., 2019; Maekawa and Fairn, 2015), we observed an increase in lysosome cholesterol levels in senescent cells **(Fig. 1 E,F and Fig. S1 A)**, which may underpin the insensitivity of mTORC1 activity to exogenous cholesterol feeding.

**Figure 1.**
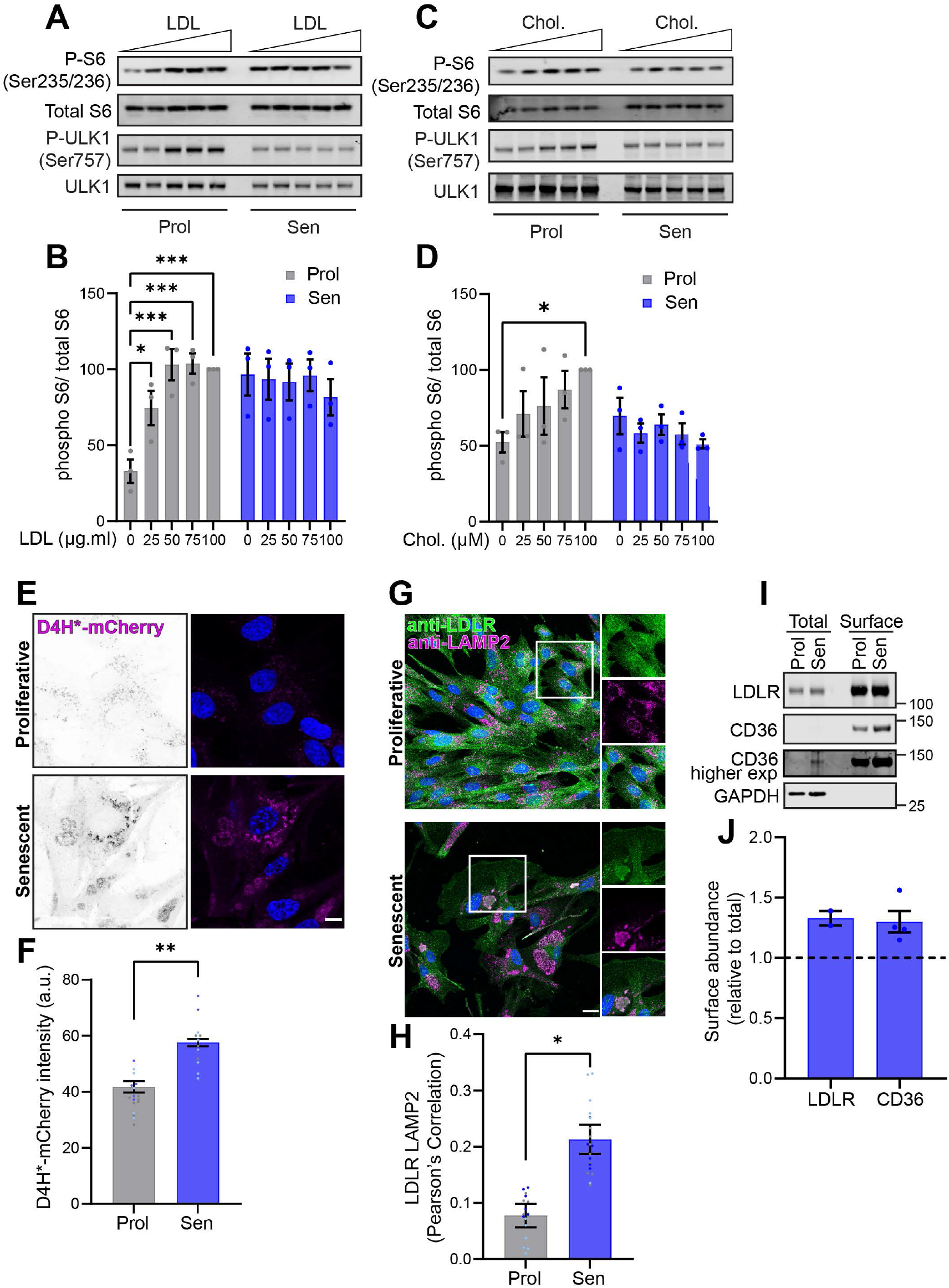
Cholesterol-dependent mTORC1 activation is perturbed in senescence. (A) Representative immunoblot analysis of dose-dependent activation of mTORC1 by LDL in proliferative or senescent cells. IMR90 ER:RasV12 primary fibroblasts were cultured in the presence of EtOH (Prol; proliferative) or 100 nM 4OHT for 6–8 days to induce RasV12 expression and oncogene-induced senescence (Sen; senescent). Cells were depleted of sterol with methyl-β cyclodextrin (MCD, 0.5% w/v) for 2 hours and then stimulated for 2 hours with increasing concentrations (0 to 100 μg/ml) of low-density lipoprotein (LDL). Cell lysates were analysed for phosphorylation of S6 (Ser235/236) and ULK1 (Ser757). (B) Quantification represents mean ± SEM; n = 3 independent experiments; two-way ANOVA followed by Dunnett’s multiple comparison. (C) Representative immunoblot analysis of activation of mTORC1 signalling by cholesterol. Proliferative or senescent cells were treated as stated in (A), however stimulated for 2 hours with increasing concentrations (0 to 100 μM) of MCD:cholesterol. (D) Phosphorylation of S6 (Ser235/236) was quantified as in (B). (E) Proliferating or senescent cells were fixed, permeabilised using a liquid-nitrogen pulse and subjected to cholesterol labelling by GST-D4H*–mCherry. Scale bar, 10 μm. The quantification of D4H*–mCherry signal intensity is shown in (F). The quantification represents the mean intensity ± SEM compared by Student’s t test (unpaired); n = 3 independent experiments with at least 5 fields of view per experimental repeat. (G) Proliferative or senescent cells were fixed and immunostained for antibodies against LDLR (green) and LAMP2 (magenta), scale bar, 20 μm. (H) Colocalisation analysis of endogenous LDLR and LAMP2. The graph represents means ± SEM. Pearson’s correlation; n = 3 independent experiments with ≥ 6 fields of view per experimental repeat. Student’s t test (unpaired). (I) Proliferating and senescent cells were surface biotinylated, and streptavidin agarose was used to capture biotinylated membrane proteins. Surface abundance of the indicated proteins was assessed by immunoblotting. (J) The quantification shows the mean ± SEM surface levels in senescent cells presented relative to proliferating cells (dashed line); n = 2 independent experiments for LDLR n = 4 independent experiments for CD36. *, P < 0.05; **, P < 0.01; ***, P < 0.001.

Cholesterol is delivered to the lysosome via the LDL receptor (LDLR), which, when bound to cholesterol-laden LDL is endocytosed and transported to the lysosome where cholesterol can drive mTORC1 activation. Increased colocalisation of LDLR with the lysosomal marker LAMP2 in senescent cells could suggest cholesterol accumulation is the result of increased delivery **(Fig. 1 G, H)**. Interestingly, despite the increase in lysosomal LDLR, cell surface biotinylation revealed the receptor is still abundant at the cell surface in senescent cells **(Fig. 1. I, J)**. Consistently, the cell surface receptor, CD36, which among other roles is a fatty acid translocase and contributes to cholesterol homeostasis, is also elevated at the cell surface of senescent cells. Together, these results indicate that mTORC1 is insensitive to exogenous cholesterol feeding in senescence, which may be mediated by increased uptake, delivery and accumulation in the lysosome. These results are interesting because one would expect delivery of LDLR to the lysosome would result in degradation and a reduction in total protein levels, however total protein levels of LDLR remain unchanged but there is clear evidence of increase surface localisation, and a simultaneous increase in colocalisation with the lysosomal marker, LAMP2.

We and others have extensively characterised lysosomes in senescence; they have higher pH, evidence of increased damage and show reduced degradative capacity (Curnock et al., 2023). These data may suggest that a LAMP2-positive organelle in senescence is vastly different to one in a proliferating cell. This would have a major but completely unexplored impact on upstream process, including trafficking through the endo-lysosomal network. Reanalysis of our proteomics dataset of isolated lysosomes (Curnock et al., 2023) identified that lysosomes from senescent cells were abundant in non-lysosomal proteins including the endocytic markers, Rab5c and the FYVE-domain containing Rab5 effector Early Endosome Antigen 1 (EEA1). In addition to Rab5c, the Rab GTPases, Rab14 and Rab8, were also detected in senescent cell lysosomes (Curnock et al., 2023). Consistently, Rab5 and EEA1 were enriched on LAMP2-positive structures, along with the phosphoinositide, PI(3)P probe GFP-2xFYVE in senescence and Rab5-LAMP2 colocalisation was significantly increased compared to proliferating controls **(Fig. 2 A-D)**. Proliferating cells however, showed clear distinction between endosomes and lysosomes with GFP-2xFYVE, Rab5 and EEA1 present on the same puncta which were absent for the lysosomal marker LAMP2 **(Fig. 2 A-D)**. These data suggest there is an accumulation of hybrid endo-lysosomal compartments in senescence.

**Figure 2.**
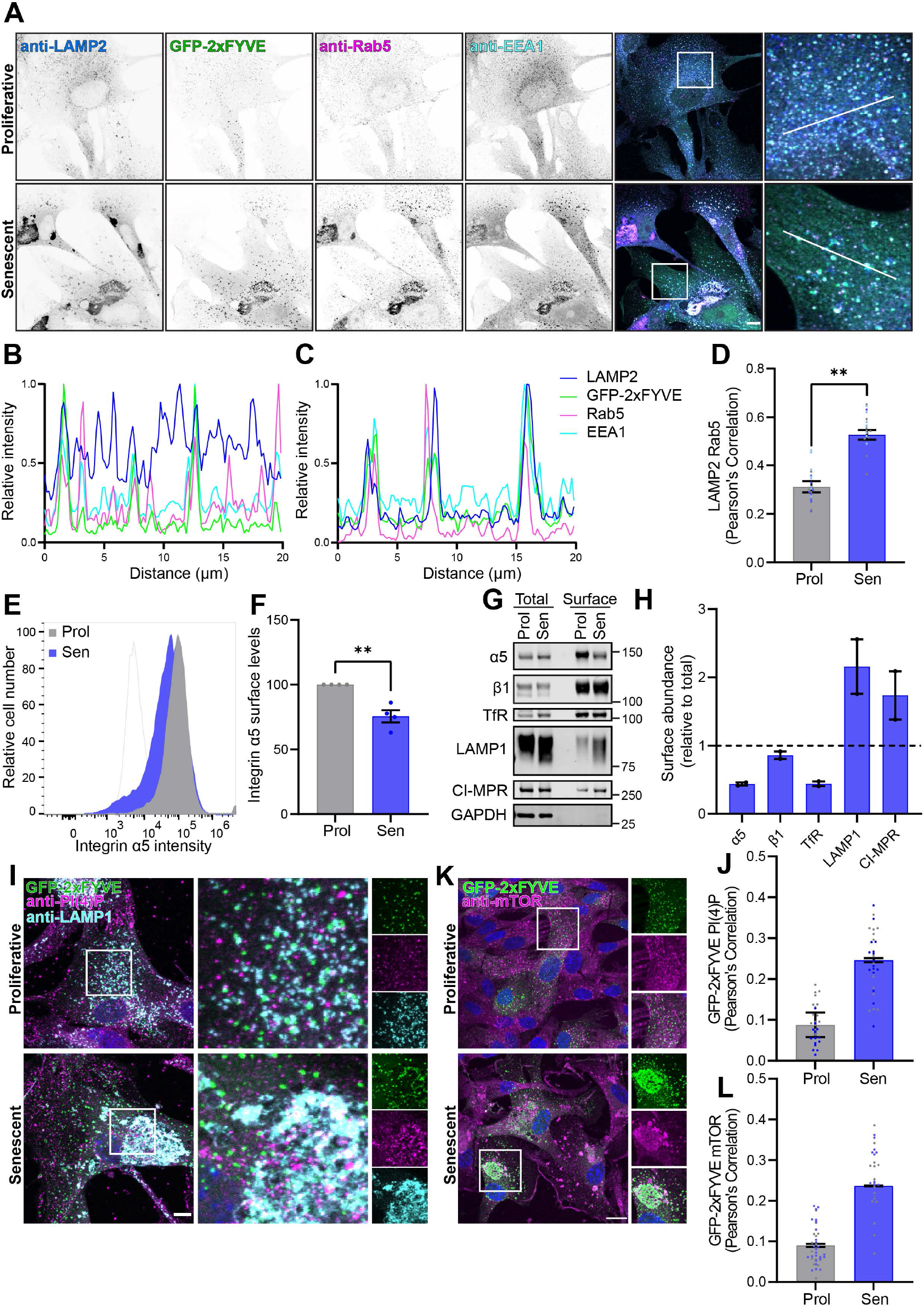
The senescence-associated hybrid endo-lysosomal phenotype causes cargo protein missorting. (A) IMR90 ER:RasV12 primary fibroblasts expressing GFP-2xFYVE were cultured in the presence of EtOH (Prol; proliferative) or 100 nM 4OHT for 6–8 days to induce RasV12 expression and oncogene-induced senescence (Sen; senescent). Cells were fixed and immunostained for antibodies against LAMP2 (blue), Rab5 (magenta) and EEA1 (cyan), scale bar, 10 μm. The signal intensity profile of the line in the merged zoom panels are shown in (B) for the proliferative and (C) for the senescent cell. Colocalisation between LAMP2 and Rab5 (D) was analysed by Pearson’s correlation. Error bars represent SEM; n = 3 independent experiments with at least 7 fields of view per experimental repeat. Student’s t test (unpaired). (E) Cell surface integrin α5 levels were analysed by flow cytometry in proliferative and senescent cells. (F) Quantification represents the change in integrin α5 cell surface levels in senescent cells versus proliferative cells. The graph represents MFI ± SEM; n = 4 independent experiments. Student’s t test (unpaired). (G) Proliferative and senescent cells were surface biotinylated and streptavidin–agarose used to capture biotinylated membrane proteins. Surface abundance of the indicated proteins were assessed by immunoblotting. (H) Quantification represents mean ± SEM surface levels in senescent cells presented relative to proliferating cells (dashed line); n = 2 independent experiments. (I) Cells were treated as stated in (A), fixed and immmunostained for antibodies against PI(4)P (magenta) and LAMP1 (cyan), scale bar 10 μm. Colocalisation between GFP-2xFYVE and PI(4)P (J) was analysed by Pearson’s correlation. Error bars represent SEM; n = 2 independent experiments with at least 14 cells in each condition per experimental repeat. (K) Proliferating and senescent cells expressing GFP-2xFYVE were fixed and immunostained with an antibody against mTOR (magenta), scale bar 20 μm. (L) The colocalisation between GFP-2xFYVE and mTOR in proliferating and senescent cells was analysed by Pearson’s correlation. Quantification represents mean ± SEM; n = 2 independent experiments with at least 16 cells per experimental repeat. **, P < 0.01.

To determine the implications of these changes on the endosomal system we assessed the total and surface abundance of well-characterised endocytic cargo proteins, integrin α5β1 and the transferrin receptor. Surface expression of integrin α5 was reduced in senescent cells, as assessed by flow cytometry **(Fig. 2 E, F)**. Cell surface biotinylation confirmed this loss was surface specific and not due to a reduction in total integrin α5 protein levels **(Fig. 2 G, H)**. Supporting these findings, we also previously identified integrin α5 to be abundant in lysosomal preps from senescent cells (Curnock et al., 2023). Relative to their total protein abundance, integrin β1 and the transferrin receptor were also lost from the cell surface in senescent cells **(Fig. 2 G and H)**. In contrast, the lysosomal proteins, LAMP1 and CI-MPR, were enriched in cell surface fractions from senescent cells **(Fig. 2 G, H)** which is consistent with previous reports that there may be enhanced lysosomal exocytosis in senescence (Bae et al., 2022; Rovira et al., 2022). Further evidence for the existence of hybrid lysosomal compartments in senescence is the increased co-existence of organelles enriched with both the phosphoinositides, PI(3)P (marked by GFP-2xFYVE) and PI(4)P (labelled with primary antibodies) **(Fig. 2 I, J)**. Membrane phosphoinositide species tightly controls the endo-lysosomal pathway and lysosomal function. PI(3)P and PI(4)P, have recently been shown to mark motile mTORC1-active and static degradative lysosomes, respectively (Ebner et al., 2023). In proliferating cells, we found that the PI(3)P probe GFP-2xFYVE was dispersed throughout the cell from the perinuclear region to the periphery and displayed restricted colocalisation with mTOR **(Fig. 2 K, L)**, whereas in senescent cells we found that mTOR was readily recruited to PI(3)P-positive (denoted by GFP-2xFYVE) structures **(Fig. 2 K, L)**.

Lysosomal damage has previously been demonstrated to simultaneously increase in PI(4)P and lysosomal cholesterol but inhibit mTORC1 (Ebner et al., 2023; Tan and Finkel, 2022). However, in senescent cells expressing the PI(4)P marker GFP-P4M-SidMx2 we observed that mTOR was recruited to cholesterol- and PI(4)P-positive structures **(Fig. S2 A, B)**, providing further evidence that senescent cells possess atypical lysosomes **(Fig. 2 I, J)**. Together, these data indicate that there is a ‘loss of identity’ of lysosomes in senescence, and instead there is an increase in hybrid organelles enriched in markers for endosomal and lysosomal compartments and the accumulation of mistrafficked proteins.

We previously showed that lysosomal pH is higher in senescence (Curnock et al., 2023) and thus we propose that the mistrafficked cargo is not degraded efficiently, and ultimately accumulates leading to the observed lysosomal membrane damage (Curnock et al., 2023; Johmura et al., 2021). In response to lysosomal damage, the kinase, phosphatidylinositol-4 kinase type 2α (PI4K2A), has recently been shown to accumulate at the lysosomal membrane to generate PI(4)P and coordinate the transfer of cholesterol from endoplasmic reticulum (ER) to the lysosome, thus restoring lysosomal membrane integrity (Radulovic et al., 2022; Tan and Finkel, 2022). Given that senescent lysosomes are characterised by increased membrane damage and increased PI(4)P, we hypothesised that this PI4K2A-dependent pathway may be upregulated in an attempt to repair lysosomes. In support of this, in senescent cells we observed an increase in colocalisation between PI4K2A and LAMP1 **(Fig. 3 A, B)**. The presence of galectin-3 positive puncta is commonly used to identify elevated lysosomal membrane damage (Papadopoulos and Meyer, 2017). We found that PI4K2A-positive puncta intersected on LysoTracker-positive galectin-3 puncta **(Fig. S2 C-E)**.

**Figure 3.**
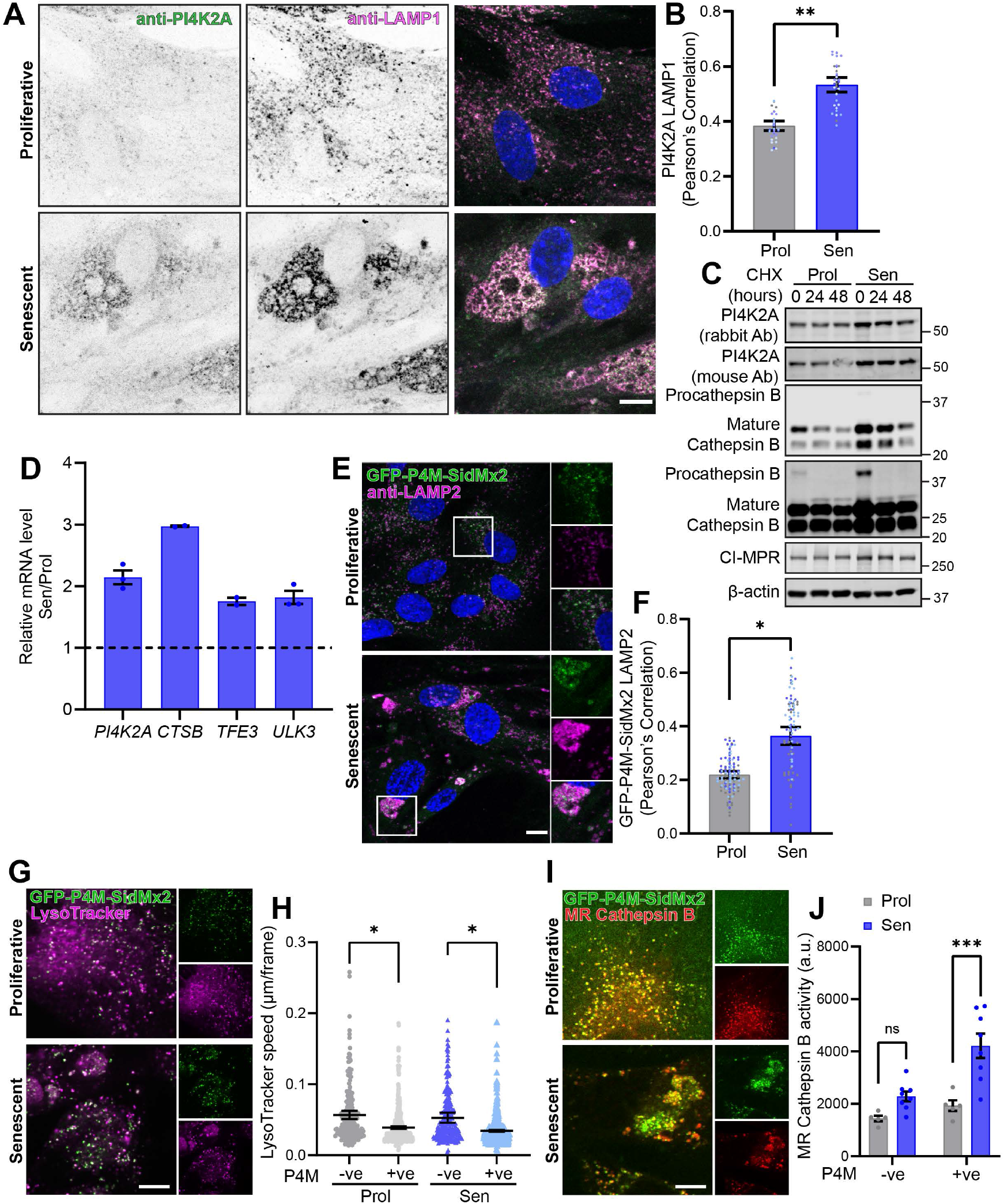
PI4K2A is upregulated in senescence. (A) Proliferative and senescent cells were fixed and immunostained for antibodies against PI4K2A (green) and LAMP1 (magenta), scale bar 10 μm. Colocalisation between PI4K2A and LAMP1 (B) was analysed by Pearson’s correlation. Error bars represent SEM; n = 5 independent experiments with at least 6 fields of view per experimental repeat. Student’s t test (unpaired). (C) Proliferative and senescent cells were incubated with 50 μg/ml cycloheximide (CHX) for the timepoints indicated. The abundance of PI4K2A and the indicated proteins were assessed by western blot. (D) mRNA levels of senescent cells, normalised to *PUM1* and presented relative to proliferating cells (dashed line) n = 3 independent experimental repeats for *PI4K2A* and *ULK3* and n = 2 independent experimental repeats for *CTSB* and *TFE3*. (E) Representative immunofluorescence image of proliferative and senescent cells expressing the PI(4)P sensor GFP-P4M-SidMx2 immunostained for an antibody against LAMP2 (magenta), scale bar 10 μm. (F) Colocalisation between GFP-P4M-SidMx2 and LAMP2 was analysed by Pearson’s correlation. Error bars represent SEM; n = 3 independent experiments with at least 32 cells per experimental repeat. Student’s t test (unpaired). (G) Representative movie stills of live-cell imaging experiment showing LysoTracker-labelled lysosomes (magenta) in proliferating and senescent cells expressing the PI(4)P sensor GFP-P4M-SidMx2. Scale bar, 10 μm (H) The speed of PI(4)-positive and -negative LysoTracker-labelled lysosomes (μm/frame) are displayed. The mean ± SEM speed from 6 movies per condition are compared by two-way ANOVA followed by Šídák’s multiple comparison. (I) PI(4)P-positive lysosomes in senescent cells display higher MagicRed Cathepsin B activity compared to PI(4)P-positive lysosomes in proliferating cells. Representative movie stills of live-cell imaging experiment showing MagicRed Cathepsin B-labelled lysosomes (red) in proliferating and senescent cells expressing the PI(4)P sensor GFP-P4M-SidMx2. Scale bar, 10 μm. (J) Quantification of PI(4)P-positive and -negative MagicRed Cathepsin B activity in proliferating or senescent cells. Quantification represents mean ± SEM; n = 2 with ≥ 5 movies analysed per condition; two-way ANOVA followed by Šídák’s multiple comparison. *, P < 0.05; **, P < 0.01; ***, P < 0.001;

The PI4K2A-dependent lysosomal repair pathway was first described in response to acute lysosomal damage (Radulovic et al., 2022; Tan and Finkel, 2022). Following cycloheximide treatment, we observed sustained elevated PI4K2A protein levels in senescent cells compared to proliferating cells **(Fig. 3 C)**, suggesting this pathway can also be employed for long-term lysosomal maintenance. Senescence is characterised by transcriptional upregulation of lysosomal quality control via TFEB (**Fig. 3 D**, (Carroll et al., 2017; Curnock et al., 2023), and here we show that this extends to *PI4K2A*. There was a 2-fold increase in *PI4K2A* mRNA levels in senescence cells compared to proliferating cells **(Fig. 3 D)**, and consistently, a significant increase in PI(4)P levels (visualised by GFP-fused PI4P-binding domain from the bacterial effector protein SidM (Hammond et al., 2014)) on senescent lysosomes **(Fig. 3 E, F)**. Live cell imaging of cells expressing GFP-P4M-SidMx2 and labelled with LysoTracker or MagicRed Cathepsin B confirmed, as previously reported that PI(4)P-positive lysosomes mark less motile and more proteolytically active lysosomes (Ebner et al., 2023) in both proliferating and senescent cells **(Fig. 3G-J and S3 A-D)**. However, in senescent cells we found that degradative (MagicRed Cathepsin B-positive) lysosomes are less mobile **(Fig. S3 D)** and more proteolytically active **(Fig. 3 I, J and S3 C)** than those in proliferating cells **(Fig. 3 G-J and S3 A-D)**, akin to lysosomes that accumulate in starved cells (Ebner et al., 2023).

The PI4K2A-dependent lysosomal repair pathway relies on membrane contacts between the endoplasmic reticulum (ER) localised VAP-A/B proteins and the cholesterol-PI(4)P transporter oxysterol-binding protein (OSBP) at the lysosome. In response to damage, OSBP facilitates the transport of PI4P and cholesterol between ER and damaged lysosome, this exchange enhances lysosomal cholesterol content which provides membrane stability (Radulovic et al., 2022; Tan and Finkel, 2022). We therefore sought to determine the localisation of OSBP to assess whether this lysosomal repair machinery is initiated in senescent cells. In proliferating cells, consistent with previous reports, GFP-OSBP was predominantly in a perinuclear Golgi region **(Fig. 4 A)**. Similar to observations in lysosomotropic agent treated cells, the GFP-OSBP signal in senescent cells was more dispersed with increased puncta that colocalised with LAMP2 **(Fig. 4 A and B)**. Importantly, in agreement with the OSBP and VAP-A lysosomal repair machinery being intact, immunoprecipitation confirmed the interaction between OSBP and VAP-A was unaltered in senescence **(Fig. S4 A)**. Interestingly, if enhanced lysosomal PI(4)P in senescence **(Fig. 3 E, F)** is supporting ER-to-lysosome transport of cholesterol (Goto et al., 2016; Mesmin et al., 2017; Mesmin et al., 2013), this could underpin or contribute to the observed increase in lysosomal cholesterol **(Fig. 1 E, F and S1 A)**. Together, these data suggest that senescent cells engage the so-called phosphoinositide-initiated membrane tethering and lipid transport (PITT) pathway, recruiting PI4K2A, enhancing lysosomal PI(4)P and cholesterol, presumably in an attempt to repair lysosomal damage and restore lysosomal function. We and others have shown that transcriptional upregulation of lysosomes is important for supporting mTORC1 activity and senescent cell survival (Curnock et al., 2023), and we hypothesized that the PITT pathway could provide further support.

**Figure 4.**
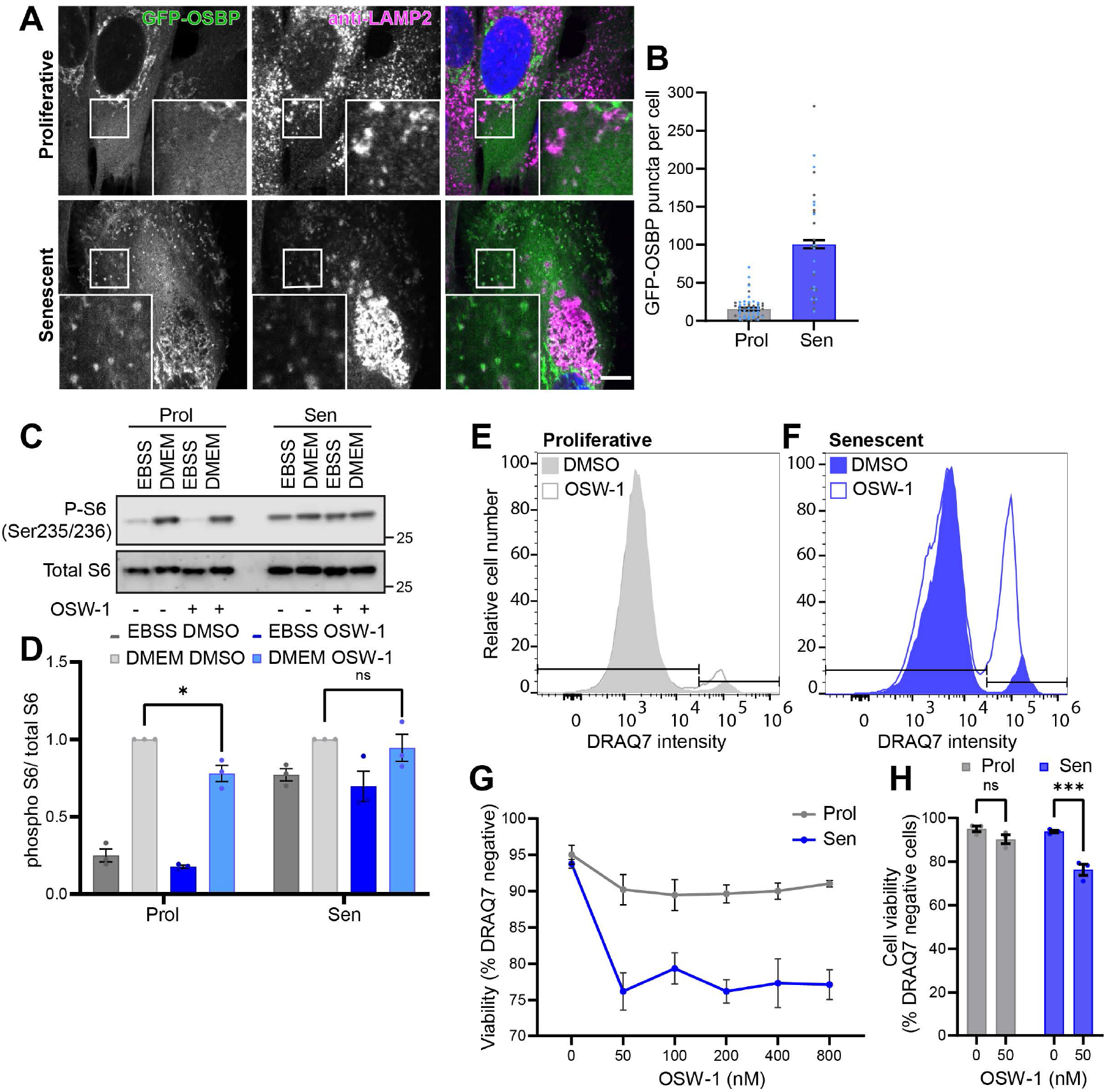
The phosphoinositide-initiated membrane tethering and lipid transport (PITT) pathway is required for senescent cell viability. (A) IMR90 ER:RasV12 primary fibroblasts expressing GFP-OSBP were cultured in the presence of EtOH (Prol; proliferative) or 100 nM 4OHT for 6–8 days to induce RasV12 expression and oncogene-induced senescence (Sen; senescent). Cells were fixed and immunostained with an antibody against LAMP2 (magenta), scale bar, 10 μm. (B) The number of GFP-OSBP puncta (defined as 0.5-0.05 μm^3^) per cell were plotted as mean ± SEM, n = 2 independent experiments with at least 14 cells per experimental repeat. (C) Proliferating and senescent cells were serum starved overnight, then treated with DMSO or OSW-1 for 4 hours and incubated with EBSS or stimulated with complete media (DMEM) for the last hour of treatment. Cell lysate were analysed for phosphorylation of S6 (Ser235/236) (D) Quantification represents mean ± SEM; n = 3 independent experiments; two-way ANOVA followed by Dunnett’s multiple comparison. (E) and (F) DRAQ7 intensity measured by flow cytometry reported the death of proliferating (E) and senescent (F) following 4 hours treatment with DMSO or 50 nM OSW-1. (G) Proliferating and senescent cell viability was measured (% DRAQ7 negative cells) following 4 hours treatment with increasing concentrations (0 to 800 nM) of OSW-1. (H) Cell viability after 4 hours treatment with 50 nM OSW-1 was analysed by two-way ANOVA followed by Šídák’s multiple comparison. Quantification represents mean ± SEM; n = 3 independent experiments. *, P < 0.05; ***, P < 0.001.

The PITT pathway can be disrupted by inhibition of OSBP with OSW-1, which leads to an accumulation of PI(4)P and has been linked to reduced mTORC1 lysosomal recruitment and activity, as well as cell death upon lysosomal damage (Ebner et al., 2023; Lim et al., 2019; Radulovic et al., 2022). Inhibition of OSBP has been shown to markedly impair basal and cholesterol-induced mTORC1 activation and also modestly affect acute mTORC1 activation by amino acids (Ebner et al., 2023; Lim et al., 2019). In nutrient depleted conditions inhibition of OSBP with acute OSW-1 treatment however did not restore any mTORC1 sensitivity in senescence **(Fig 4 E, F)**. Whereas, consistent with previous observations (Lim et al., 2019), in proliferating cells, treatment with OSW-1 caused a modest reduction in mTORC1 activity in response to acute stimulation by nutrients **(Fig 4 E, F)**. Despite the sustained elevated lysosomal recruitment of PI4K2A and concomitant increase in lysosomal PI(4)P, mTORC1 activity persists in senescent cells even in the absence of nutrients or lipids **(Fig 4 E, F)**. Together, this suggests that senescent cells are unresponsive to nutrient-regulated PI(3)P and PI(4)P functional switching at the lysosomal membrane and mTORC1 activation is driven autonomously to the supply of exogenous nutrients.

Importantly however, OSW-1 promoted an increase in cell death in senescent cells **(Fig. 4 G-J)**, but had no effect on cell viability, even at higher concentrations in proliferating cells **(Fig. 4 I)**. We found that OSW-1 induced senescent cell death in the same timeframe to that observed upon induction of lysosomal damage in OSBP depleted cells (Radulovic et al., 2022), indicating that cell survival after acute or sustained lysosomal membrane damage relies on OSBP-dependent PI(4)P and cholesterol transfer. Furthermore, these data suggests targeting lysosomal repair mechanisms has senolytic potential.

In summary, we have identified that the discrete protein and phosphoinositide heterogeneity that exists in healthy endosome and lysosome populations is lost in senescence. Instead they possess a senescence-associated hybrid endo-lysosomal phenotype with the endosomal GTPase Rab5 and PI(3)P-binding protein EEA1 coexisting on LAMP2-positive lysosomes, which may be contributing to the mislocalisation of surface proteins to the lysosome. Senescent cells also fail to induce the lysosomal PI(4)P-PI(3)P phosphoinositide switch to tune lysosomal function and mTORC1 activity (Ebner et al., 2023). Instead, these lipids converge on morphologically and functionally hybrid lysosomal populations; they possess both PI(3)P and PI(4)P and are proteolytically active yet augment mTOR recruitment. Our data indicates that sustained lysosomal damage in senescence is driving PI4K2A recruitment and the simultaneous increase in PI(4)P and lysosomal cholesterol. Perturbation of this lysosomal membrane repair pathway causes senescent cell death.

Aging and many age-related neurodegenerative conditions are linked to altered endocytic function and complex changes throughout the autophagy-lysosomal system (Nixon and Rubinsztein, 2024; Palmer et al., 2025; Small and Petsko, 2020). These pathways all converge at the lysosome. Impaired lysosomal acidification is linked to Alzheimer’s disease and other age-related diseases (Colacurcio and Nixon, 2016). However, it is still unclear whether lysosomal dysfunction is a direct cause or consequence of defects that arise in aging, senescence, and neurodegenerative diseases. The importance of the lysosome to cellular homeostasis and ultimately cell survival is demonstrated by the evolutionarily conserved pathways that repair or degrade damaged lysosomes. Here, we show perturbation of the recently described PITT pathway may represent a novel target for senolytic development. Improving our understanding of lysosomal biology in senescence could reveal more potential senolytic targets and inventions for age-associated diseases.

## Acknowledgements

BC and RC are supported by The Royal Society and Wellcome Trust (218547/Z/19/Z awarded to BC). The authors gratefully acknowledge the Wolfson Bioimaging Facility and the Bristol Flow Cytometry Unit, University of Bristol, UK, for their support and assistance in this work.

## Figure legends

## Supplementary figure legends

S1 (A) Proliferating and senescent fibroblasts were fixed, permeabilised using a liquid-nitrogen pulse, subjected to cholesterol labelling by GST-D4H*–mCherry and immunostained with an antibody against LAMP2 (blue).

S2 (A) Proliferating and senescent cells expressing GFP-P4M-SidMx2 were fixed, permeabilised using a liquid-nitrogen pulse, subjected to cholesterol labelling by GST-D4H*–mCherry and immunostained with an antibody against mTOR (cyan). (B) Colocalisation between GFP-P4M-SidMx2 and mTOR was analysed by Pearson’s correlation. Error bars represent SEM; n = 2 independent experiments with at least 20 cells in each condition per experimental repeat. (C) Cells were incubated with 75 nM of LysoTracker Deep Red (shown in cyan) for 30 min, then fixed and immunostained for antibodies against Galectin-3 (green), PI4K2A (magenta), scale bar, 10 μm. The signal intensity profile of the line in the merged panels are shown in (D) for the proliferative and (E) for the senescent cell.

S3 (A and B) Quantification of the LysoTracker intensity (A) mean ± SEM and LysoTracker-labelled lysosomes speed (μm/frame) (B) from live-cell imaging experiment of LysoTracker-labelled lysosomes in proliferating and senescent cells with 6 time-lapse sequences compared by Student’s t test (unpaired) mean ± SEM per condition. (C and D) Quantification of and MR Cathepsin B activity (a.u.) (C) and MR Cathepsin B speed (μm/frame) (D) from live-cell imaging experiment of MagicRed Cathepsin B-labelled lysosomes comparing by Student’s t test (unpaired) mean ± SEM of 5 proliferating and 8 senescent time-lapse sequences.

S4 (A) Western blot of GFP immunoprecipitation from proliferating and senescent cells expressing GFP or GFP-OSBP.

## Materials and methods

### Cell culture and drug treatments

The IMR90 cells (human primary fibroblasts) stably expressing inducible HRas^V12^ were cultured in phenol red-free DMEM supplemented with 2 mM glutamine, 20% FBS, 1 mM sodium pyruvate, 1× non-essential amino acids and 100 U/ml penicillin–streptomycin at 37°C, 5% CO^2^ and 3% O^2^. To induce OIS, HRasV12 expression was induced by treatment of cells with 100 nM hydroxy-tamoxifen (Sigma-Aldrich, H7904) for 6–8 days. OSW-1 was obtained from Cayman Chemical (30310), and used at concentrations from 50 nM to 800 nM for 4 hours. Cells were incubated with 50 μg/ml cycloheximide (CHX) for the indicated time. CHX was obtained from Sigma-Aldrich (C1988).

### DNA plasmids

pGEX-KG-D4H*-mCherry (#134604) and the lentiviral vectors; pLJM1-FLAG-GFP-OSBP (#134659), pLenti-EGFP-P4M-SidMx2 (#136997) and pLenti-EGFP-2xFYVE (#136996) were all obtained from Addgene.

### Viral production

All proteins were expressed through lentiviral infection. Lentiviruses were produced by co-transfecting HEK293T cells with the viral vector containing the gene of interest and packaging plasmids pAX2 and pMDG2 using polyethylenimine (PEI, 47336, Thermo Fisher Scientific) transfection method. Viral supernatant was collected 72 hours after transfection, filtered with a 0.45 μm filter and incubated with fibroblasts overnight. Media was changed the following day and cells cultured for the indicated amount of time.

### Cell lysis and immunoblotting

Cells were rinsed with cold PBS and lysed in lysis buffer (RIPA buffer; 50 mM Tris–HCl, pH 7.4, 150 mM NaCl, 1% NP-40, 0.5% sodium deoxycholate, and 0.1% SDS, supplemented with Halt protease and phosphatase inhibitors, PI78443, Thermo Fisher Scientific) on ice. Cell lysates were cleared by centrifugation at 17,000 g for 10 minutes at 4 °C, and protein concentration was measured using DC protein assay (500-0112; Bio-Rad Laboratories), and equal amounts of protein were resolved on NuPAGE 4–12% gradient gels. Proteins were transferred from gels onto PVDF membrane (Immobilon-FL, pore size 0.45 μm, Millipore) for immunoblotting. Transfer was performed in transfer buffer (25 mM Tris, 192 mM glycine and 10% methanol) at 100 V for 70 min. Membranes were blocked in 5% milk in PBS containing 0.1% Tween 20 (PBS-T) prior to incubation with the appropriate primary antibodies and fluorescently labelled secondary antibodies. Blots were imaged on LiCOR Odyssey xL (LiCOR) and quantified using ImageLite software (LiCOR).

The following primary antibodies were used: mouse monoclonal antibodies raised against Lamp1 (H4A3, DSHB, 1:2000), PI4K2A (sc-390026, Santa Cruz, 1:2000), β-Actin (8H10D10, 3700, Cell Signalling Technology, 1:5000), GAPDH (D4C6R, 97166, Cell Signalling Technology, 1:2000), integrin beta 1/CD29 (610467, BD Biosciences, 1:2000), transferrin receptor/CD71 (3B8 2A1, sc-32272, Santa Cruz, 1:1000), GFP (7.1/13.1, Roche, 11814460001, 1:2000), rabbit monoclonal antibodies raised against cathepsin B (D1C7Y, 31718, Cell Signalling Technology, 1:2000), CI-MPR (EPR6599, 124767, Abcam, 1:2000), Lamp1 (D2D11 9091, Cell Signalling Technology, 1:2000), phospho-S6 Ser235/236 (4856, Cell Signaling Technology, 1:2000), S6 (2217, Cell Signaling Technology, 1:1000), phospho-ULK1 Ser757 (D7O6U, 14202, Cell Signaling Technology, 1:1000), ULK1 (D8H5, 8054, Cell Signaling Technology, 1:1000), integrin alpha 5 (EPR7854, ab150361, Abcam, 1:2000), rabbit polyclonal antibodies raised against PI4K2A (15318-1-AP, Proteintech, 1:1000), LDLR (10785-1-AP, Proteintech, 1:2000), CD36 (18836-1-AP, Proteintech, 1:1000), VAPA (15275-1-AP, Proteintech, 1:2000). Secondary antibodies conjugated to Alexa Fluor 680 or 800 were used at 1:20000.

### GFP immunoprecipitation

For immunoprecipitation experiments, cells expressing GFP-tagged proteins were placed on ice, the medium was removed and the cells were washed twice times with ice-cold PBS (Sigma). Cells were lysed in ice-cold Tris-based immunoprecipitation buffer (50 mM Tris-HCl, pH 7.4, 0.5% NP-40 supplemented with Halt protease and phosphatase inhibitors; 1861280; Thermo Fisher Scientific). And the lysates were cleared by centrifugation. Fifteen microlitres of GFP-trap beads (Chromotek, catalogue number gta-20) were equilibrated prior to adding cell lysate supernatants. Beads and lysates were incubated on a rocker at 4 °C for 1 hour. Following incubation, GFP-trap beads were pelleted and the supernatant was removed. Beads were then washed by resuspending in lysis buffer, pelleting and removal of lysis buffer. This washing process was repeated three times in total. Once the final lysis buffer wash had been removed, beads were resuspended in 2× NuPAGE LDS Sample Buffer (Life Technologies), 2.5% β-mercaptoethanol and boiled at 95 °C for 10 minutes. The samples were separated by SDS-PAGE and subjected to immunoblot analysis.

### Cell surface biotinylation

Cells were surface biotinylated with membrane-impermeable biotin (21331, Thermo Fisher Scientific) at 4°C to prevent endocytosis. Following biotinylation, cells were lysed (2% Triton X-100, PBS, supplemented with Halt protease and phosphatase inhibitors; 1861280; Thermo Fisher Scientific, pH 7.5), and lysates were cleared by centrifugation. Equal amounts of protein from the control and indicated knockout were then added to streptavidin Sepharose to capture biotinylated proteins. Streptavidin beads and lysates were incubated for 30 min at 4°C before washing in PBS containing 1.2 M NaCl and 1% Triton X-100. Proteins were eluted in 2× NuPAGE LDS Sample Buffer (Life Technologies) by boiling at 95°C for 10 minutes and then were separated by SDS-PAGE and subjected to immunoblot analysis.

### Cholesterol starvation and stimulation

IMR90 cells were rinsed twice with serum-free media and incubated in 0.5% w/v methyl-beta cyclodextrin (MCD, C4555, Sigma-Aldrich) for 2 hours. Cells were then transferred to serum-free media containing 0.1% MCD (starved condition) or 0.1% MCD supplemented with 25-100 μg/ml low-density lipoprotein (LDL, 437644, Sigma-Aldrich) or 25-100 μM cholesterol (C3045, Sigma-Aldrich) pre-complexed with 0.1% MCD and incubated for 2 hours.

### Immunofluorescence

Cells grown on coverslips were fixed in 4% formaldehyde/PBS for 15 minutes at room temperature. Cells were then washed in PBS, permeabilised with 0.1% Triton X-100/PBS for 10 minutes, washed in PBS and blocked in 1% BSA/PBS for 30 minutes. Primary antibodies were diluted in 1% BSA/PBS and incubated for 1 h at room temperature. Primary antibodies used in this study: mouse monoclonal antibodies raised against PI4K2A (sc-390026, Santa Cruz, 1:500), Lamp1 (H4A3, DSHB, 1:2000), Lamp2 (H4B4, DSHB, 1:2000), PI(4)P (z-p004, Echelon, 1:100), rabbit monoclonal antibody raised against Rab5 (C8B1, 3547, Cell Signalling Technology, 1:200), Lamp1 (D2D11 9091, Cell Signalling Technology, 1:500), mTOR (7C10, 2983, Cell Signaling Technology), rabbit polyclonal antibodies raised against LDLR (10785-1-AP, Proteintech, 1:400), Galectin-3 (14979-1-AP, Proteintech, 1:200) and sheep polyclonal antibody raised against EEA1(AF8047, R&D Systems, 1:400).

Cells were washed in PBS and incubated with the appropriate Alexa Fluor-conjugated secondary antibodies (1:500; Thermo Fisher Scientific) in 1% BSA/PBS containing DAPI for 30 minutes at room temperature. Coverslips were mounted on glass slides using Fluoromount-G. Where indicated cells were incubated with 75 nM of LysoTracker Deep Red 30 minutes, then fixed in 4% formaldehyde/PBS and immunostained as above for Galectin-3 and PI4K2A. Confocal images were collected on Leica SP5-II AOBS confocal laser scanning microscope using 63× HCX Plan-Apo/1.3-NA oil objective. Images were analysed with the Volocity 6.3.1 software (PerkinElmer). For colocalisation studies, Pearson’s correlation (measuring the correlation in the variation between two channels) were measured using manual threshold method.

Endogenous PI(4)P staining has been described previously (Hammond et al., 2009). Briefly, cells grown on coverslips were washed once with PBS, fixed for 15 minutes at room temperature with 2% PFA, 2% sucrose/PBS, washed three times with 50 mM NH4Cl/PBS, permeabilised for 5 minutes at room temperature with 20 μM digitonin (14952, Cayman Chemical) diluted in buffer A (20 mM Pipes, pH 6.8, 137 mM NaCl, 2.7 mM KCl), washed three times with buffer A, blocked for 1 hour at room temperature with 1% BSA in PBS/50 mM NH4Cl. Cells were then washed twice in buffer A, incubated for 45 minutes at room temperature with primary antibodies against PI(4)P (z-p004, Echelon, 1:100) and Lamp1 (D2D11 #9091, Cell Signalling Technology, 1:500) diluted in 1% BSA/PBS, washed three times in PBS, incubated for 1 hour room temperature with the appropriate Alexa Fluor-conjugated secondary antibodies (1:500, Thermo Fisher Scientific) in 1% BSA/PBS containing DAPI, washed three times in PBS, post-fixed in 2% PFA, 2% sucrose/PBS, washed in 50 mM NH4Cl/PBS and mounted on glass slides using Fluoromount-G.

### Labelling of cholesterol with GST–D4H*–mCherry

Purification of recombinant GST–D4H*–mCherry and labelling of cholesterol was performed as previously described (Lim et al., 2019). Briefly, cells grown on coverslips were fixed in 4% PFA for 10 minutes at room temperature and permeabilised in a liquid nitrogen bath for 30 seconds. Cells were then blocked in 1% BSA/PBS for 1 hour and incubated in recombinant GST–D4H*–mCherry diluted in 1% BSA/PBS for 2 hours. Cells were then washed twice in PBS, fixed for a second time in 4% PFA for 10 minutes, and where indicated immunostained for Lamp2 (H4B4, DSHB, 1:2000) or mTOR (7C10, 2983, Cell Signaling Technology), coverslips were washed in PBS and mounted on glass slides using Fluoromount-G.

### Live-cell imaging of PI(4)P and LysoTracker or MagicRed Cathepsin B

Cells expressing the PI(4)P fluorescent reporter GFP-P4M-SidMx2 were seeded on 35 mm glass-bottom dishes (MatTek), and 48-72 h later, cells were incubated with either 75 nM of LysoTracker Deep Red (L12492, Thermo Fisher Scientific) for 5 minutes or Magic Red Cathepsin B (ICT937, Bio-Rad) for 1 hour and then imaged using an Olympus IXplore SpinSR system at 37 °C in 5% CO2. For tracking of lysosomes (LysoTracker or MagicRed Cathepsin B), 1 frame per second time-lapse sequences were recorded for 120 seconds. Speed and intensity measurements were performed using a custom-made ImageJ (Fiji) plugin.

### RNA extraction, reverse transcription, and quantitative PCR

Total RNA was extracted from proliferative or senescent IMR90 cells using RNeasy Mini Kit (Qiagen). cDNA was synthesised and real-time quantitative RT–PCR was performed using the SuperScript III Platinum SYBR Green One-Step quantitative RT–PCR Kit (Thermo Fisher Scientific). The quantification of gene expression was performed in triplicate. Amplification of the sequence of interest was normalised with a housekeeping gene, PUM1. Quantification was performed using comparative threshold cycle (Ct) method of analysis. The value was expressed as a fold change relative to RNA from proliferating cells.

### Flow cytometry

Cells were incubated in complete media containing 0 to 800 nM of OSW-1 for 4 hours. Cells were washed in PBS (without Ca2+ or Mg2+) and then incubated for 20 minutes with Accutase to detach cells. Resuspended cells were subsequently transferred to a 5 ml round-bottomed polystyrene tubes (BD Biosciences). Cells were reconstituted in 100 μl CO2-independent media supplemented with 5% FCS and DRAQ7 (DR71000, Biostatus, 1:100). DRAQ7 intensity from at least 20,000 cells were measured using ACEA Novocyte 3000 analyser. Data were analysed using Flowjo software (Flowjo, Ashland, OR).

### Statistical analysis

Statistical analyses were performed using Prism 7 (GraphPad Software). For comparing two groups, statistical analyses were performed using Student’s *t* test (unpaired and two-tailed). For multiple comparisons the following statistical tests were used: two-way ANOVA followed by Šídák’s multiple comparison or Dunnett’s multiple comparison. Graphs represent means and SEM.

## Notes

### Competing Interest Statement

The authors have declared no competing interest.

