## Supplementary figures and images for "Senescent cell survival relies on upregulation of lysosomal quality control mechanisms"

### Supplementary Figures 1-4

#### Supplementary figure 1

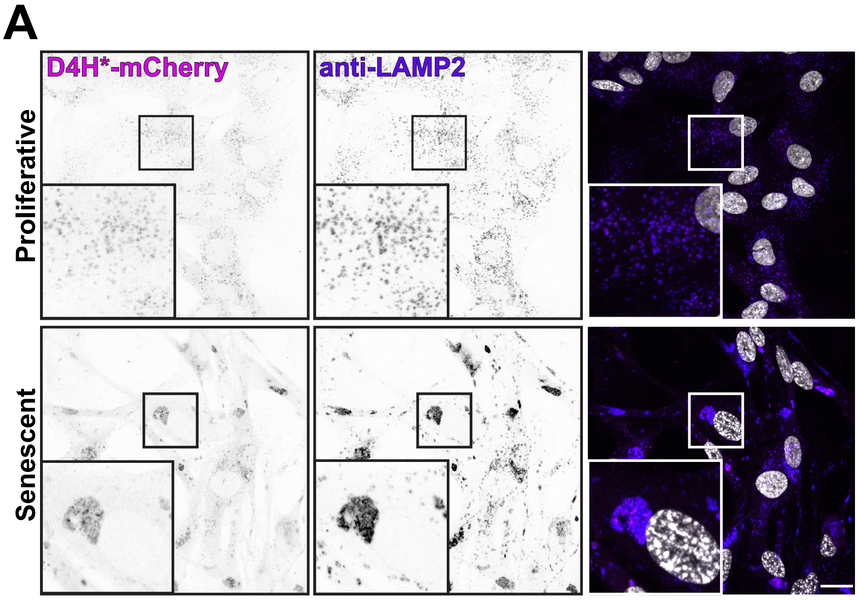

Supplementary figure 2

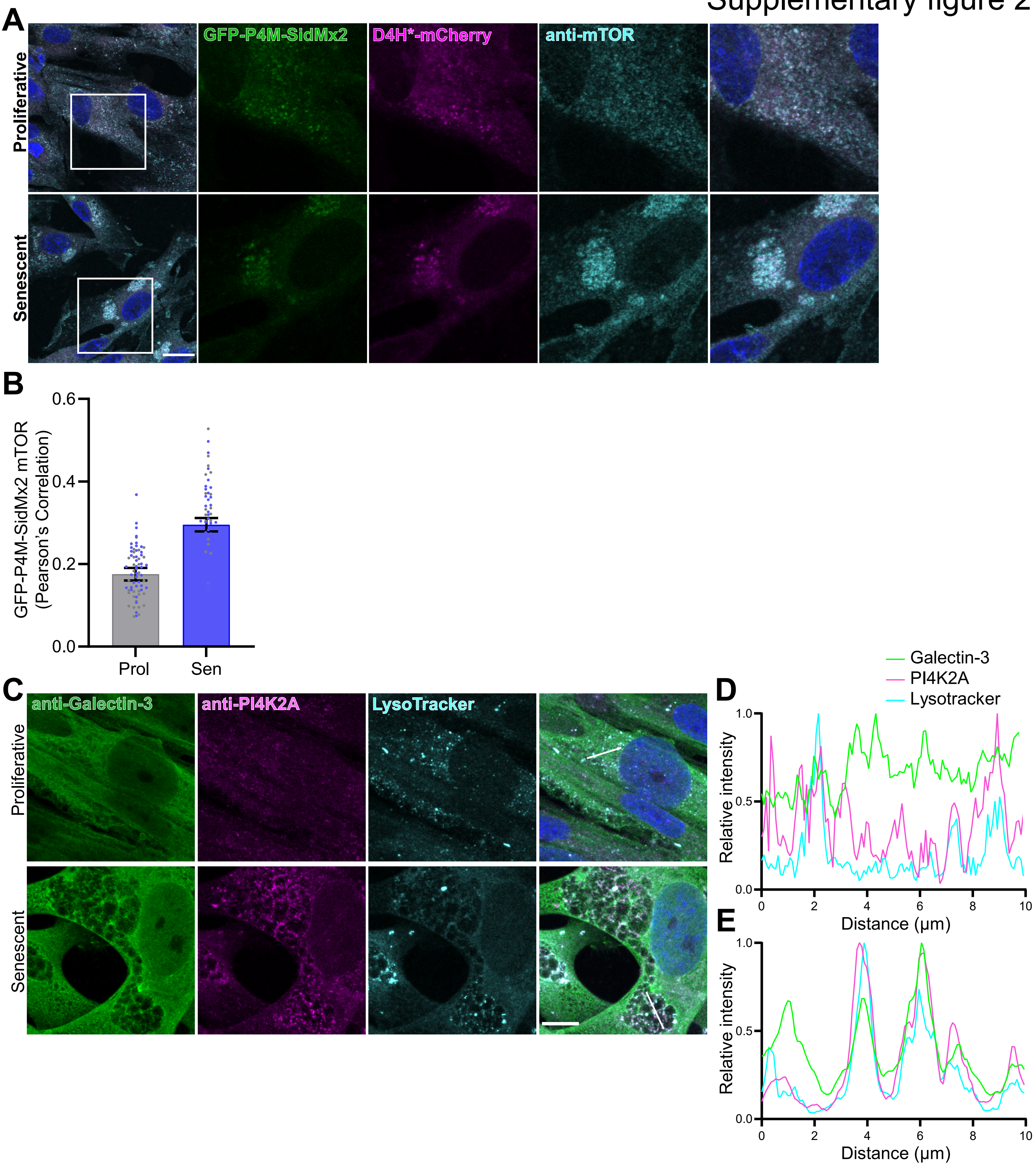

#### Supplementary figure 3

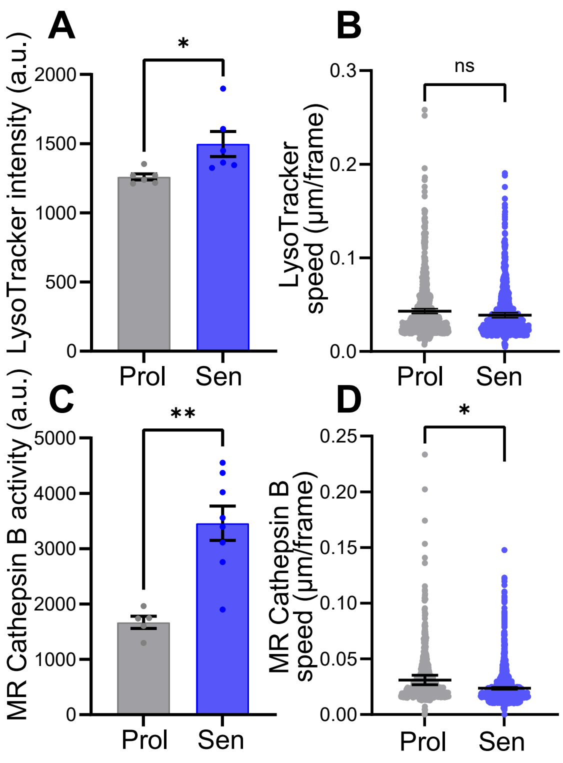

Supplementary figure 4

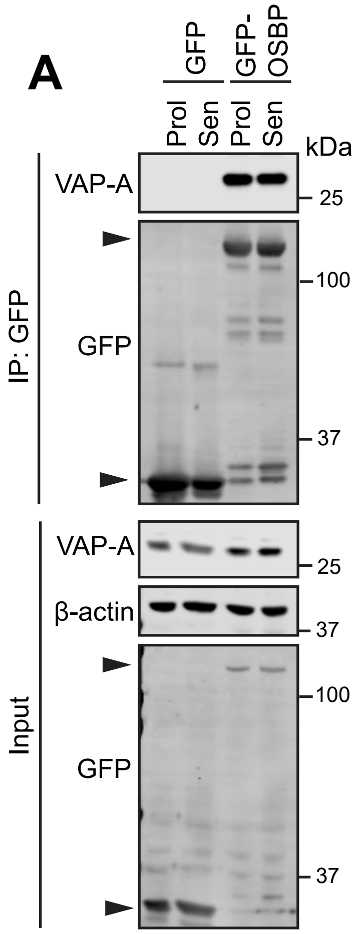
